# Diverse chemical functionalization of nucleobases within long RNAs using sulfinate salts

**DOI:** 10.1101/2021.10.19.465008

**Authors:** Anastassia Gomez, Tiziano Bassi, Leah Grayson, Julien Vantourout, Navtej Toor

## Abstract

We have devised a single pot, low-cost method to modify RNA with sulfinate salts that can directly add almost any desired functional group to nucleobases under mild aqueous conditions. This chemistry modifies the Hoogsteen edge of RNA and DNA nucleobases. It can be applied to RNA or DNA of any size, as well as to individual nucleotides. Existing methods of RNA modification have relatively limited applicability due to constraints on the size of the RNA and the lack of diversity of possible modifications. We have been able to add azide groups for click reactions directly onto the nucleobases of RNA utilizing sulfinate salts. C-H bonds on the nucleobase aromatic rings serve as the sites of attachment, with C-H being replaced with C-R, where R is the azide-containing linker. With the addition of azide functional groups, the modified RNA can easily be reacted with any alkyne-labeled compound of interest, including fluorescent dyes as shown in this work. This methodology enables the exploration of diverse chemical groups on RNA that can potentially confer protection from nucleases, allow for efficient delivery of nucleic acids into cells, or act as new tools for the investigation of nucleic acid structure and function.

## Introduction

There are currently over 170 different RNA modifications that are known to exist in prokaryotes and eukaryotes^1^. These RNA modifications affect all aspects of gene expression and regulation, including transcription and translation. These modifications occur on the nucleobase aromatic rings and are incorporated through the action of specific enzymes for each functional group that is added to the RNA^1^. One of the most prevalent modifications is the presence of pseudouridine in mRNA. This modification greatly increases the halflife of pseudouridine-containing RNA *in vivo* compared to unmodified RNA^2^. Remarkably, pseudouridine is able to stabilize RNA *in vivo* with a random distribution of this residue throughout the mRNA sequence. Pseudouridine and its derivatives are now the basis for the multi-billion dollar industry that develops mRNA therapeutics and produces vaccines for COVID-19^3^. mRNA is also modified with other functional groups in the cell, such as N6-methyladenosine, to regulate its translation by the ribosome^4^. A recent report of acetyl modifications to cytidine nucleobases in mRNA showed that these modifications promoted translation efficiency and mRNA stability *in vivo*^5^. Therefore, RNA modification is an emerging area of study that is revealing itself to be a significant mechanism for modulating biological pathways and conferring new properties upon RNA.

Given the impact of nucleobase modifications upon the biochemical properties of RNA, there has been great interest in the incorporation of novel synthetic functional groups to enable additional functionality. Reagents such as dimethyl sulfate and kethoxal have been historically used to modify the Watson-Crick edge of nucleobases in RNA for structure probing^6^; however, it is not possible to add a wide variety of functional groups using this type of chemistry. The synthesis and chemical diversity of synthetically modified RNA is limited by the following problems:

1. T7 RNA polymerase does not efficiently incorporate modified nucleobases during *in vitro* transcription. Long chemically modified RNAs (>100 nucleotides) are typically synthesized using T7 RNA polymerase in the presence of modified nucleoside triphosphates (NTPs). However, T7 RNA polymerase is not able to accommodate highly modified and/or bulky unnatural NTPs within the restrictive confines of its active site^7,8^. As a result, the diversity of modified RNA is limited by the enzymatic mechanism of T7 RNA polymerase. The majority of possible modifications cannot be incorporated at all by the polymerase.
2. The low efficiency of incorporation of modified NTPs during *in vitro* transcription results in exorbitant costs of synthesis. Modified NTPs are extremely expensive and the low efficiency of incorporation greatly increases the cost of synthesis for large-scale production of therapeutics^8^.
3. Chemical synthesis to incorporate modified nucleobases is largely limited to the production of short nucleic acids. Solid phase oligonucleotide (oligo) synthesis is limited to the modification of small nucleic acids; generally, no more than 200 deoxyribonucleotides with the phosphoramidite method and no more than 60 ribonucleotides for RNA^9^. For example, direct synthesis of modified messenger RNAs (typically >1,000 nucleotides) is not currently possible using chemical methods. A method to bioconjugate DNA oligonucleotides was recently published but the technology requires a solid support and is limited to short DNAs^10^.

RNA poses unique challenges for chemical modification due to its relative lability compared to DNA^2,11,12^. The modification of RNA has generated interest in medicinal chemistry due to its potential use as a therapeutic agent for a range of diseases^2,11,13–15^. Unfortunately, many reactions used in synthetic organic chemistry would result in the RNA being destroyed due to the harsh conditions. The modification of nucleic acids is also broadly useful as a research tool by the addition of bioconjugation tags for affinity purification or by the addition of fluorophores for cellular tracking of labeled RNAs^11,12,16^. Further development of RNA-based therapeutics would greatly benefit from a low-cost and simple method of introducing desired functional groups to RNA nucleobases. For example, mRNA-based vaccines require the rapid production of billions of doses in a short period of time. This vaccine could be engineered with novel functional groups to exhibit greater stability within the human body for a stronger immune response to the encoded antigen. This necessitates the development of chemistry that could easily scale to the large quantities of synthetic mRNA required for vaccination, along with a low cost for the incorporation of modified nucleotides.

Here we demonstrate the chemical modification of RNA with different sulfinate salt reagents that attach various R groups to the Hoogsteen edge of the nucleobases (Figures 1A and 1B). Specifically, we tested zinc trifluoromethanesulfinate (TFMS-Zn), zinc bis(phenylsulfonylmethanesulfinate) (PSMS-Zn), and sodium (difluoroalkylazido)sulfinate (DAAS-Na). Each reagent adds a unique functional group onto the substrate. TFMS-Zn adds a trifluoromethyl group, PSMS-Zn adds a phenylsulfonylmethyl group, and DAAS-Na adds a difluoroalkylazido group in which the terminal azide moiety serves as a click chemistry substrate^17–19^. In addition, the chemistry is flexible with respect to its substrates, including both small and large oligonucleotides, as well as large sequences of RNA. Importantly, the nucleic acid functionality enabled by sulfinate salt chemistry is extremely diverse, considering the number of sulfinate salt reagents and the vast number of commercially available “clickable” moieties readily available to scientists.

**Figure 1.**
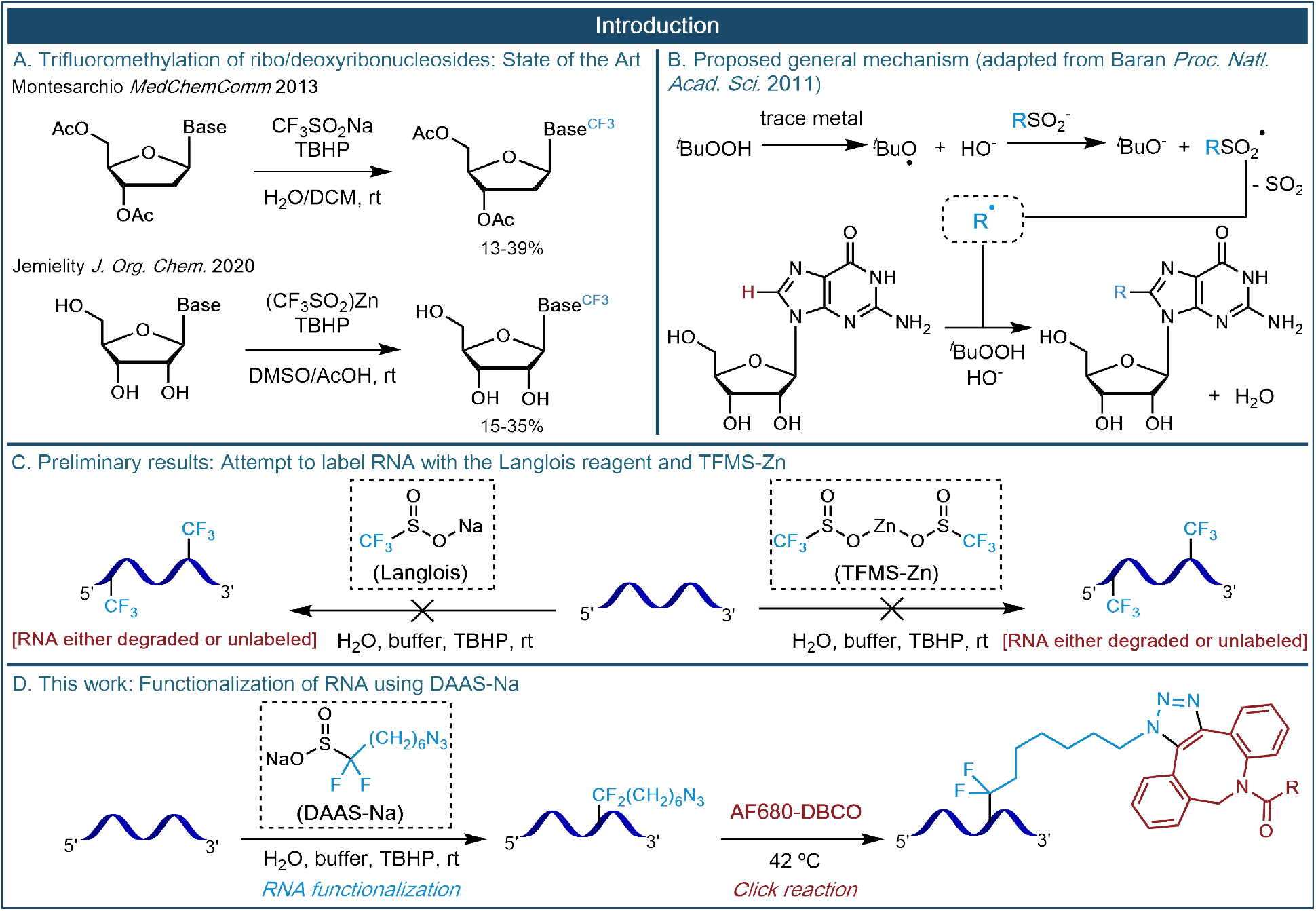
(above): A synopsis of what is currently known with regards to sulfinate modification of nucleosides is shown along with the novel method presented in this publication for long RNAs. Panel A: currently known reactions involving sulfinate labeling of ribo- and deoxyribonucleosides. Panel B: mechanism for sulfinate labeling of nucleosides. Panel C (this work): labeling long RNAs with trifluoromethanesulfinate reagents was unsuccessful and degraded the RNA. Panel D (this work): labeling long RNAs with DAAS-Na preserves phosphodiester backbone integrity and allows further click chemistry.

## Results

### Initial RNA labeling attempts with trifluoromethanesulfinate

Sulfinate salts have been used to functionalize CH groups found in heteroarene-containing small molecule compounds^17,20^. We hypothesized that these salts could be adapted to introduce functional groups onto the nucleobases of RNA since they are also heteroarenes. In contrast to small molecules, RNA is a long polymer that is prone to cleavage under harsh conditions such as high pH, high temperature, and the presence of certain divalent cations. We therefore optimized reaction conditions to keep the RNA intact during the sulfinate modification reaction, which included an essential buffer component to control the pH to prevent RNA degradation.

Before attempting to label RNA with DAAS-Na, we wanted to explore if labeling was possible with the simpler and much more well-known trifluoromethanesulfinate salts. Sodium trifluoromethanesulfinate (TFMS-Na, often referred to as the “Langlois Reagent”) was the first of the sulfinate salts shown to be capable of labeling various aromatic structures under oxidative conditions^21^. We first modified individual DNA bases to determine sites of modification by TFMS-Zn. TFMS-modified DNA bases evaluated by fluorine-19 NMR showed a single reactive position on each nucleobase (Figure 2). Notably, all these positions are on the Hoogsteen edge of the nucleobases.

**Figure 2.**
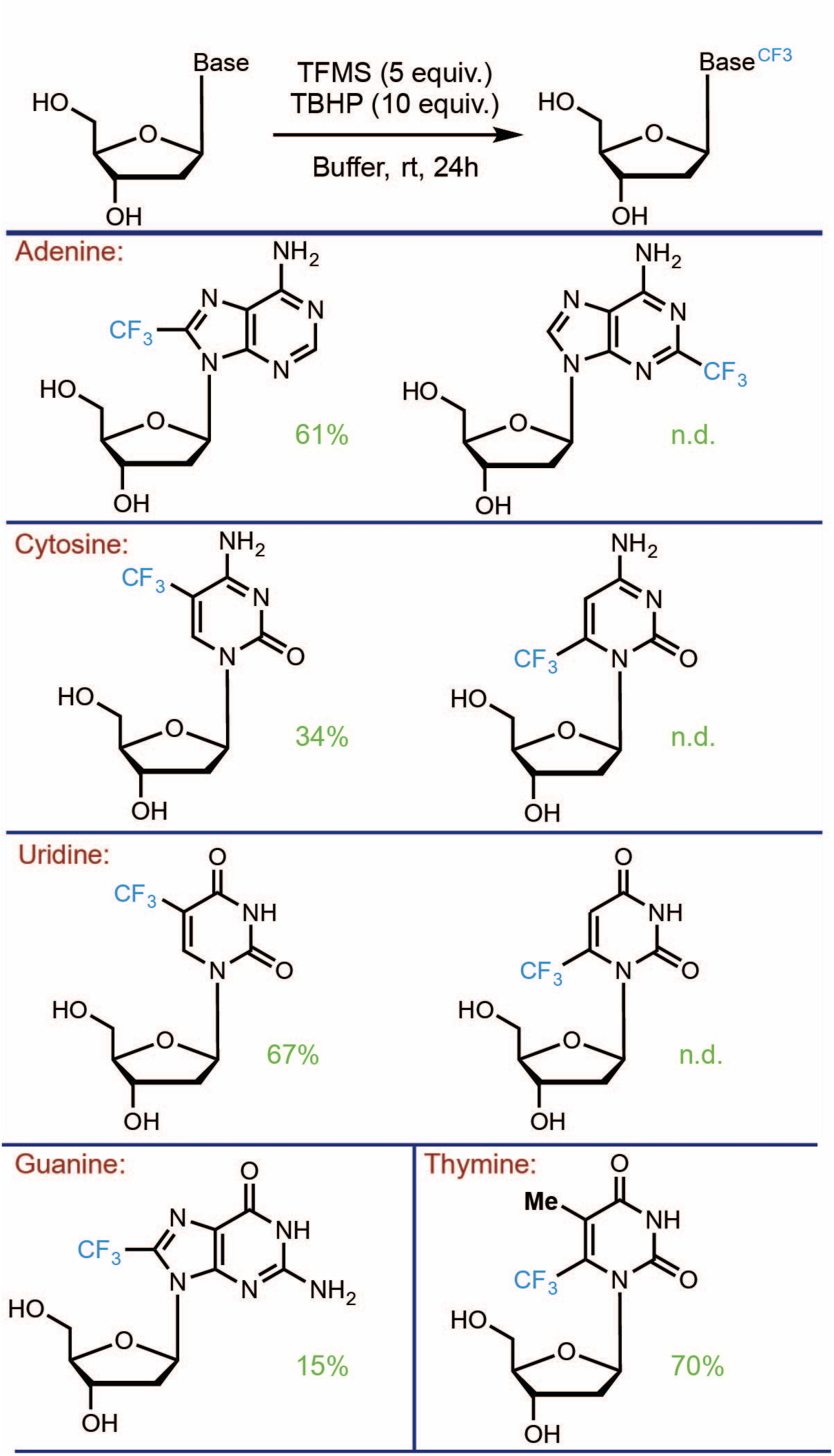
(above): Deoxyribonucleosides were modified by TFMS-Zn in separate reactions and fluorine-19 NMR was used to identify the sites of trifluoromethylation on each nucleobase. The CF_3_-thymidine ^19^F NMR yield was 70% and the ^19^F-NMR chemical shift was δ −63.1 ppm (376 MHz, CDCl_3_). The CF_3_-deoxyadenosine ^19^F NMR yield was 61% and the ^19^F-NMR chemical shift was δ −62.4 ppm (376 MHz, CDCl_3_). The CF_3_-deoxycytidine ^19^F NMR yield was 34% and the ^19^F-NMR chemical shift was δ −61.9 ppm (376 MHz, CDCl_3_). The CF_3_-deoxyguanosine ^19^F NMR yield was 15% and the ^19^F-NMR chemical shift was δ −62.1 ppm (376 MHz, CDCl_3_). The CF_3_-deoxyuridine ^19^F NMR yield was 67% and the ^19^F-NMR chemical shift was δ −62.7 ppm (376 MHz, CDCl_3_).

We then proceeded to test whether or not the RNA remained intact after being modified by TFMS-Na (Figure 1C). We found that every attempt to label RNA with TFMS-Na resulted in one of two outcomes: either (1) the RNA stayed intact but was not labeled, or (2) the RNA was successfully labeled but was heavily degraded in the process (Supplementary Figures 1 and 2; Supplementary Table 1). Labeling was verified by first enzymatically digesting the RNA into nucleoside 5’-monophosphates followed by quantitative ^19^F NMR (using a sodium trifluoroacetate internal standard). Extensive ^19^F NMR trifluoromethylation studies on pure samples of the individual nucleoside monophosphates revealed the identities of all relevant chemical shifts, showing that the four canonical RNA nucleoside monophosphates are all capable of being trifluoromethylated. Integrity of the RNA was verified with denaturing polyacrylamide gel electrophoresis (PAGE). Attempts were also made to label RNA using zinc trifluoromethanesulfinate (TFMS-Zn)^17^, but similarly to TFMS-Na, this always resulted in one of the two outcomes discussed above (Supplementary Figures 2 and 3). The factors that governed which of the two outcomes occurred were largely based on the reaction conditions, particularly the type and amount of buffer used. Large excesses of buffer relative to the sulfinate salt generally resulted in outcome (1), whereas smaller buffer concentrations generally resulted in outcome (2). This is likely due to the buffers interfering with the reaction when present at a large enough concentration. However, the question remained as to why the trifluoromethanesulfinate salts (either TFMS-Na or TFMS-Zn) destroyed the RNA under conditions in which they were allowed to react without significant buffer interference. Though multivalent transitional metal cations such as Zn^2+^ are capable of catalyzing phosphodiester bond cleavage, this cannot be the dominant factor responsible for trifluoromethanesulfinate-induced strand cleavage since TMFS-Na also causes severe RNA degradation. To ensure that trace metals in the TMFS-Na could be ruled out, we ran additional TFMS-Na labeling reactions with EDTA present, and these RNA samples were also completely degraded. We hypothesize that the main mechanism for strand cleavage may be related to the radical mechanism of the reaction. Particularly, it has been shown through kinetic analysis of selectively deuterated nucleic acids that the ribose 5’-C-H bond is particularly prone to radical abstraction, forming a 5’-carbon radical which eventually leads to phosphodiester backbone cleavage^22^. The trifluoromethyl radical intermediate in these reactions may abstract hydrogen directly from the RNA backbone or from water to form an OH’ radical which then abstracts hydrogen from the RNA backbone. Importantly, upon further attempts at labeling RNA with sulfinate salts other than TFMS-Na and TFMS-Zn, we saw that the tendency towards phosphodiester backbone degradation is highly dependent on the structure of the particular sulfinate salt used in the reaction. While labeling with TFMS-Na and TMFS-Zn may be too destructive, we saw that other sulfinate salts, such as DAAS-Na described below, are capable of labeling RNA without degradation.

### Modification of 20-nucleotide RNA and DNA oligonucleotides

We also attempted labeling 20-nucleotide DNA and RNA oligonucleotides (oligos) with PSMS-Zn and DAAS-Na to see if intact oligonucleotides could be detected after reaction with these sulfinate salts. The resulting oligos were analyzed by LC-MS to detect modification in the context of a full-length oligo (Table 1). We detected masses for the entire oligonucleotide plus the additional modification(s) from the sulfinate reactions. These experimental masses indicated the existence of 1 to 3 modifications per 20 nucleotides of DNA or RNA. Furthermore, we used click chemistry to add alkyne-biotin and alkyne-Flour 488 to DAAS-labeled DNA and RNA and found that the oligonucleotides were successfully labeled with these bulky functional groups. This data provided the first preliminary evidence that larger DNAs and RNAs could be labeled with sulfinate salts without degradation.

**Table 1.**
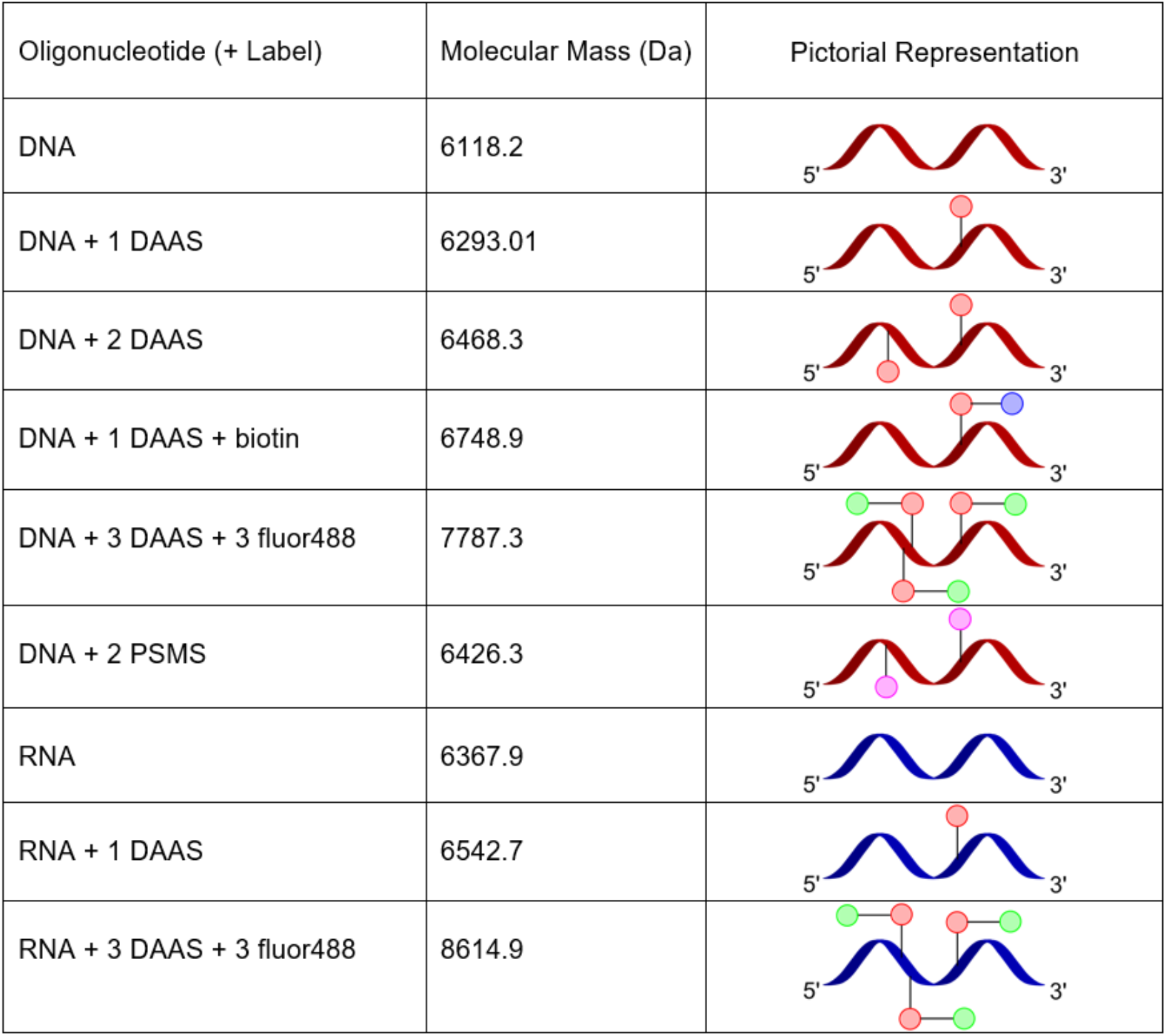
(above): LC/MS masses that were identified for various labeled DNA and RNA oligos are shown. The biotin and fluor488 labels were added via click chemistry, similarly to procedures described in this paper. Sequences are CAGAATGCTTAACGTCCGGT and CAGAAUGCUUAACGUCCGGU.

| Oligonucleotide (+ Label) | Molecular Mass (Da) | Pictorial Representation |
| --- | --- | --- |
| DNA | 6118.2 |  |
| DNA + 1 DAAS | 6293.01 |  |
| DNA + 2 DAAS | 6468.3 |  |
| DNA + 1 DAAS + biotin | 6748.9 |  |
| DNA + 3 DAAS + 3 fluor488 | 7787.3 |  |
| DNA + 2 PSMS | 6426.3 |  |
| RNA | 6367.9 |  |
| RNA + 1 DAAS | 6542.7 |  |
| RNA + 3 DAAS + 3 fluor488 | 8614.9 |  |

### Modification of a long mRNA

We then proceeded to modify an *in vitro* transcribed 1227-nucleotide (nt) GFP mRNA with DAAS-Na. Six labeling reactions (A1-A6) were prepared in aqueous solution, with concentrations of all reagents and buffers shown in Table 2. Reaction A6 was a negative control, containing all reagents except DAAS-Na. All reactions were allowed to proceed for the same length of time (overnight, approximately 23 hours) at room temperature. As a note on nomenclature, RNA that was reacted with DAAS-Na will be referred to as DAAS-RNA, indicating that azide linkers are covalently attached to the RNA. To distinguish the RNA products from different DAAS-Na labeling reactions, the reaction identifier will be appended to the end of the name (for example, the RNA labeled from reaction A3 will be referred to as DAAS-RNA A3 in future reactions that use this RNA sample for click chemistry).

**Table 2.** (above): Final concentrations of all reagents are shown for reactions A1-A6. All reactions were run in milli-Q water, at room temperature, for 23 hours.

| Labeling Reaction | GFP mRNA | Sodium Cacodylate, pH 6.5 | DAAS-Na | TBHP |
| --- | --- | --- | --- | --- |
| A1 | 1 mg/mL | 18.75 mM | 7.5 mM | 11.25 mM |
| A2 | 1 mg/mL | 25 mM | 10 mM | 15 mM |
| A3 | 1 mg/mL | 31.25 mM | 12.5 mM | 18.75 mM |
| A4 | 1 mg/mL | 37.5 mM | 15 mM | 22.5 mM |
| A5 | 1 mg/mL | 43.75 mM | 17.5 mM | 26.25 mM |
| A6 | 1 mg/mL | 25 mM | 0 mM | 15 mM |

After labeling reactions A1-A6 were complete, the DAAS-RNA was purified from the reaction mixture using molecular weight cutoff (MWCO) filtration and analyzed using denaturing polyacrylamide gel electrophoresis (PAGE), as shown in Figure 3. The presence of full-length RNA indicates that the DAAS-Na labeling reaction preserves the integrity of the RNA.

**Figure 3.**
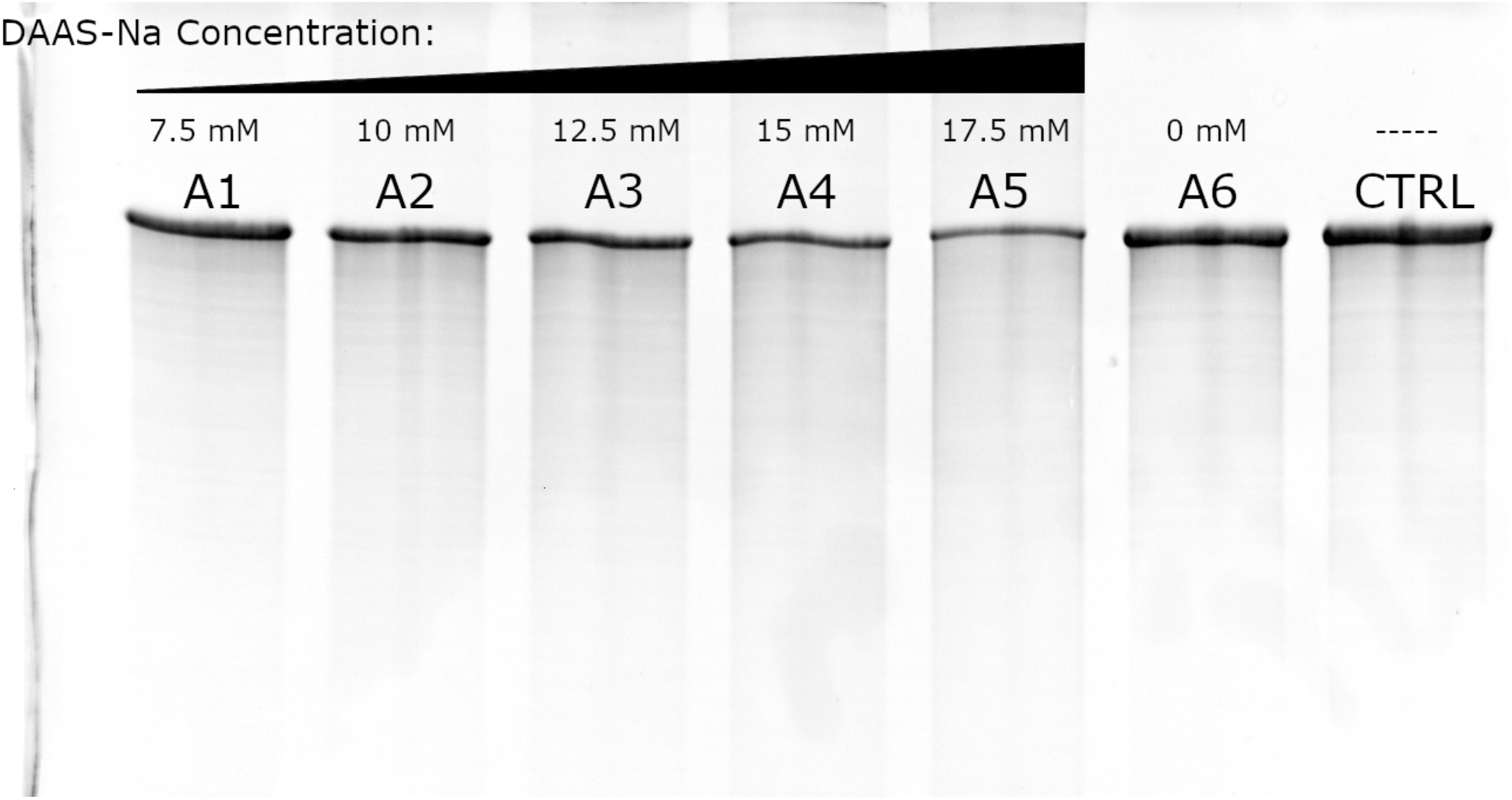
(above): Denaturing PAGE of the recovered DAAS-RNA from reactions A1-A6 (lanes 1–6) is shown along with unlabeled RNA as a control (lane 7). This gel was stained with ethidium bromide to make the RNA visible.

### DAAS-RNA can be conjugated with DBCO-labeled dye as a qualitative assay of DAAS labeling

The six DAAS-RNA samples discussed above (DAAS-RNAs A1-A6) were allowed to react at 42 °C for the same length of time (overnight, approximately 22 hours) with an excess of DBCO-labeled dye (AF680-DBCO) with the reaction scheme depicted in Figure 1D. The reaction that occurs is the well-known strain-promoted azide-alkyne cycloaddition^23^. After this, the dye-labeled RNA (referred to as DBCO-DAAS-RNA) was purified from unreacted AF680-DBCO using molecular weight cutoff filtration, with >1,000,000-fold total dilution of the original reaction mixture, ensuring no significant amounts of unconjugated dye were present with the recovered DBCO-DAAS-RNA. Though a significant blue color was noted in the flow-through of the first few MWCO filtrations (due to the AF680-DBCO being in excess), the disappearance of this color upon additional filtration cycles ensured that the amount of dye present in the final DBCO-DAAS-RNA samples was negligible. A picture of the recovered DBCO-DAAS-RNA A1-A6 samples is shown in Figure 4, with each vial on the top row containing approximately 30 μg of DBCO-DAAS-RNA and each vial on the bottom row showing the final flow-through of the MWCO filtration, indicating that there was no detectable unconjugated AF680-DBCO remaining in the DBCO-DAAS-RNA samples. UV/Vis spectroscopic analysis of the final flow-through confirmed the lack of any free dye. The labeled RNA was found to exhibit a deep blue color that was easily visible to the eye. This indicates that the level of modification of the RNA resulting in fluorophore coupling is significant enough to have large observable effect upon its biophysical properties.

**Figure 4.**
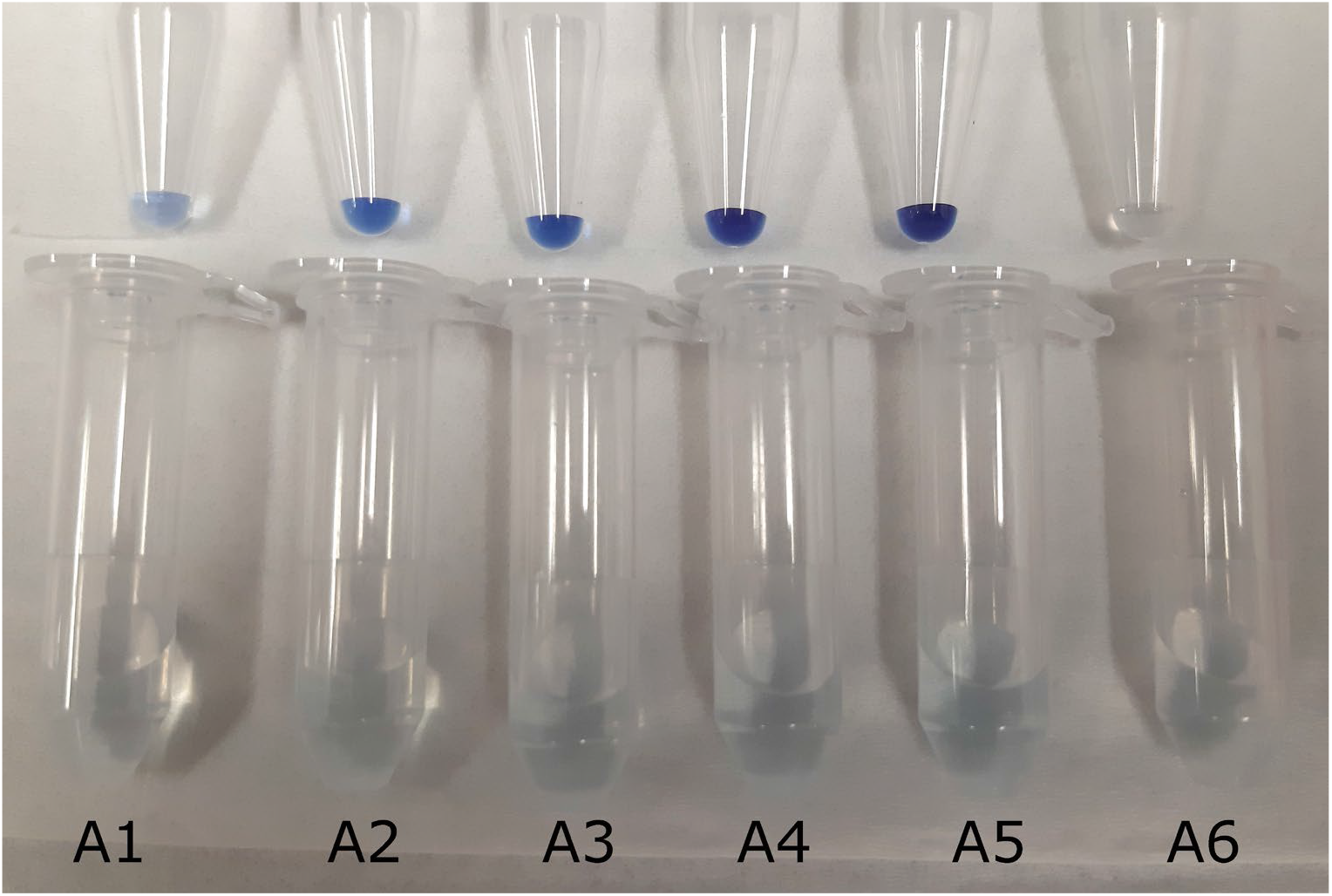
(above): Samples of DBCO-DAAS-RNAs A1-A6 are shown on the top row of tubes (approximately 30 μg per tube), with the bottom row showing the final flow-throughs of molecular weight cutoff filtration for each sample.

DBCO-DAAS-RNAs A1-A6 were run on denaturing PAGE alongside unlabeled control RNA, in the same arrangement as in Figure 3, as additional evidence of labeling. This gel was scanned at an excitation wavelength near the λ-max of the conjugated dye (~680 nm) and simultaneously imaged for fluorescence (Figure 5, Panel A). Though present on the gel, the unlabeled control RNA and RNA A6 are not visible since they do not have any covalently attached fluorophore. In contrast, DBCO-DAAS-RNAs A1-A5 gave strong signals, with the RNAs that were incubated with larger concentrations of DAAS-Na fluorescing considerably brighter, indicating higher efficiency of labeling. This gel was then ethidium bromide stained to reveal the control RNAs (Figure 5, Panel B). Taken together, these data show that DAAS-Na is capable of labeling RNA without degradation, thus allowing it to be conjugated to DBCO-labeled compounds, such as fluorescent dyes. Unexpectedly, we noted that just 2 μg of the most highly labeled sample (DBCO-DAAS-RNA A5) was able to be visually tracked by eye as it moved through the gel by simply holding a piece of white paper behind the gel apparatus and observing the blue color associated with the migrating band of DBCO-DAAS-RNA (Supplementary Figure 4).

**Figure 5.**
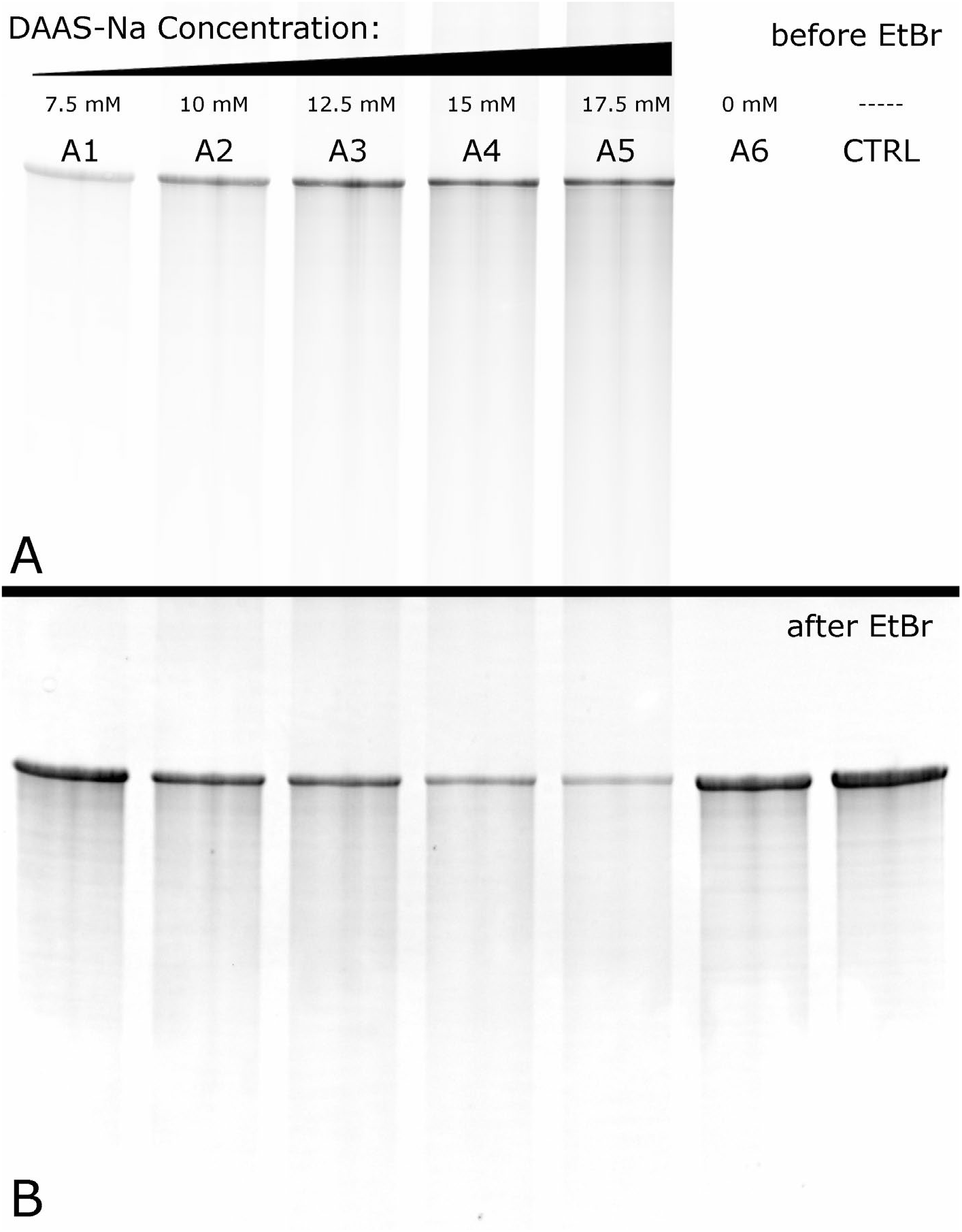
(above): Panel A: A fluorescence scan of DBCO-DAAS-RNAs A1-A6 (lanes 1–6) run on denaturing PAGE is shown, with unlabeled RNA as a control (lane 7). This gel was not yet ethidium bromide stained, hence the lack of any signal in the lane for the unlabeled RNAs. Panel B: The same gel from Panel A is shown after ethidium bromide staining. Note that the slight dimming that occurs for samples with higher labeling is not due to degradation, since both the full-mass band and streaking are proportionally diminished in intensity. Instead, we hypothesize that the increasing presence of AF680-DBCO, which absorbs in the red, is absorbing some of the red/orange emission of the ethidium bromide and is thus decreasing the signal intensity. Refer to Panel A for a better visualization of the highly labeled samples.

### The labeling percentage of DBCO-DAAS-RNA can be quantitatively assayed using UV/Vis absorbance

In order to quantitate the average number of conjugated dye molecules per DBCO-DAAS-RNA, a calibration line of 680 nm absorbance vs. AF680-DBCO concentration was made using known standards of the free dye. The calibration line and UV/Vis absorbance spectra for all four standards are shown in Supplementary Figure 5. Using this calibration line, it was possible to determine the effective concentration of AF680-DBCO dye in each sample of DBCO-DAAS-RNA. Dividing this concentration by the equivalent concentration of nucleobases within the sample yielded the average fraction of nucleobases within each sample that were conjugated with dye. This showed that up to 4% of nucleotides were labeled with fluorophore (Supplementary Table 2, Figure 6, Panel A). Also included in Figure 6 (Panel B) are UV/Vis absorbance spectra for DBCO-DAAS-RNA, unlabeled RNA, and free AF680-DBCO dye. These spectra demonstrate that DBCO-DAAS-RNA shows (1) the typical RNA peak in the 260 nm UV region from strong absorbance by the aromatic nucleobases, and (2) the characteristic 680 nm absorbance by AF680-DBCO. Unlabeled RNA looks almost identical to labeled RNA in the 260 nm UV region but fails to absorb in the visible wavelengths, while AF680-DBCO absorbs strongly in the visible region but has relatively weak absorbance in the UV. The minor absorbance of AF680-DBCO at 260 nm is convenient because this wavelength is used for RNA concentration determination, thus ensuring the fluorophore’s interference with RNA concentration is negligible.

**Figure 6.**
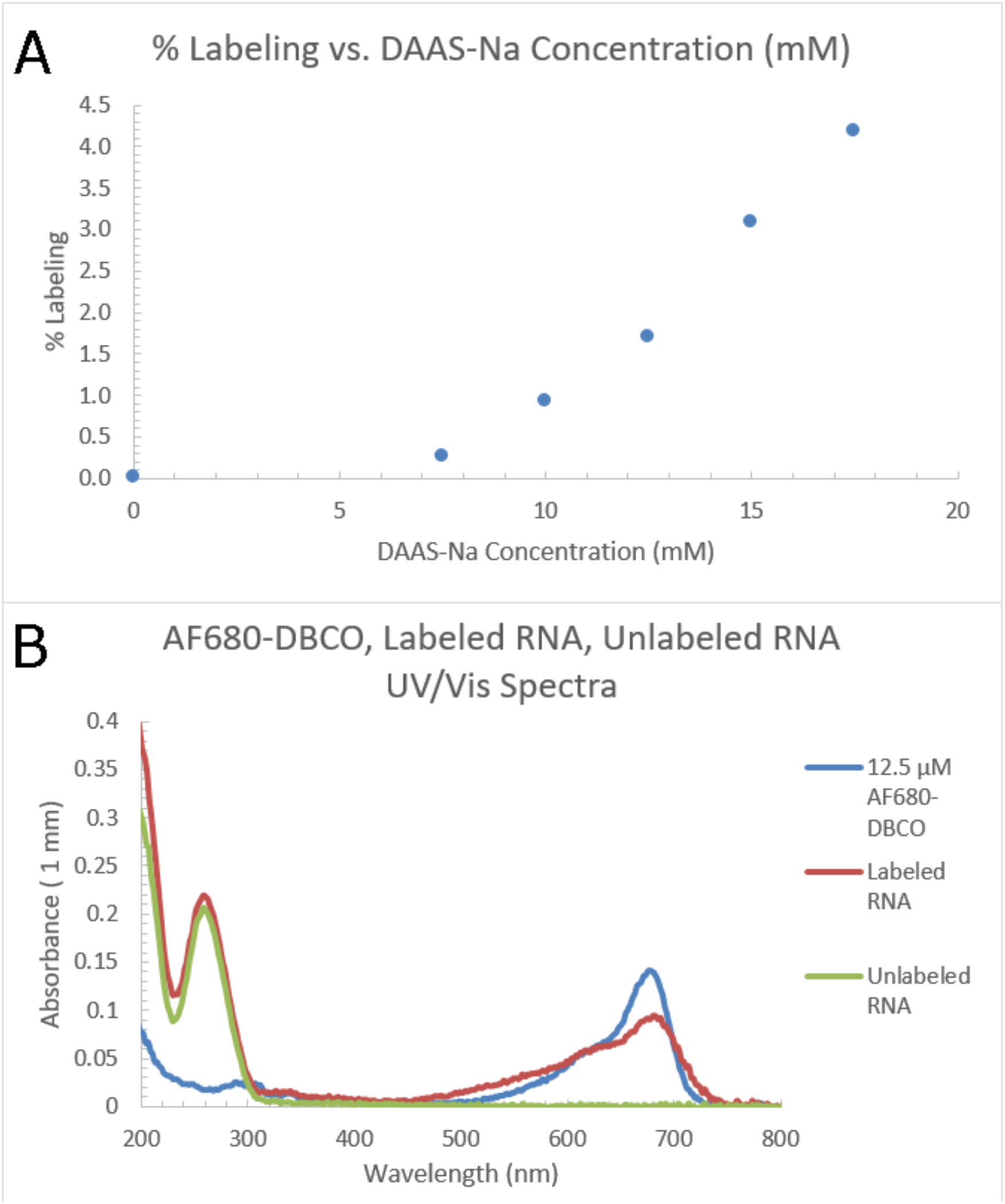
(above): Panel A: A plot of the data in Supplementary Table 4 is shown. Panel B: The UV/Vis absorbance spectra for a sample of DBCO-DAAS-RNA, a sample of unlabeled RNA, and free AF680-DBCO dye are shown superimposed onto each other.

## Discussion

We have shown a novel procedure by which azide click handles can be added to RNA without degradation through a single pot low-cost approach, allowing the RNA to be functionalized with the diverse array of commercially available DBCO small-molecule conjugates. Very large RNAs can be labeled with this method, thus allowing easy access to chemical space that otherwise would be out of reach with RNA chemical synthesis or *in vitro* transcription. Though we have shown that some sulfinate salts, such as trifluoromethanesulfinate, degrade RNA upon labeling, the ability of DAAS-Na to label RNA without significant degradation shows that sulfinate salts are capable of labeling RNA while keeping the phosphodiester backbone intact. Our mass spectrometry data confirming diverse sulfinate salt labeling of 20-nt DNA and RNA oligonucleotides provides further evidence that other sulfinate salts are suitable. These data also show that single-stranded DNA is capable of being labeled, which is expected, considering that three of the four DNA nucleobases are identical to RNA, with the fourth differing only slightly in structure. Future investigation into the chemical mechanisms behind TFMS-induced backbone cleavage will assist the search for additional sulfinate salts that are capable of labeling nucleic acids without significant degradation, increasing the diversity of functional groups that can be added.

In this work, all reactive positions are on the Hoogsteen edge of nucleic acids, which we predict enables nucleic acids to still engage in Watson-Crick pairing. This would be consistent with naturally occurring chemically modified bases that retain Watson-Crick pairing potential such as pseudouridine, which is found in eukaryotic mRNAs and is also a Hoogsteen edge modification. Modification of nucleic acids with sulfinate salts has the potential to aid development of therapeutics through the identification of moieties that can more effectively protect and efficiently deliver therapeutic nucleic acids. The random incorporation of modifications, as observed in our sulfinate protocol, is desired for some applications such as mRNA stability. Some RNA modifications, such as randomly distributed pseudouridine and N4-acetylcytidine residues, have been shown to extend the half-life of mRNA in mammalian cells. In this regard, our sulfinate methodology enables the exploration of an entire universe of functional groups that could potentially surpass the stability provided by these natural modifications. The simplicity of making modifications to RNA using sulfinate salt chemistry enables a low-cost, efficient search for modifications that could further extend the half-life of RNA *in vivo.* Similarly, this method facilitates the search for modifications that could allow RNA to more readily cross cellular membranes for the delivery of nucleic acid therapeutics. For example, covalent lipid-RNA conjugates could be generated by reacting the DAAS-modified RNA with an alkyne-labeled lipid that would result in greater hydrophobicity to enable cell entry. Optimal lipid groups could be screened quickly given the fact that there are a large number of commercially available alkyne-labeled lipids that could be used in screening experiments for efficient long RNA delivery.

Sulfinate salt chemistry is also useful as a tool for basic research of nucleic acid biochemistry. Fluorescent probe attachment can be helpful for tracking nucleic acids in gels and for localization within cells. The attachment of affinity probes, such as biotin, can facilitate nucleic acid purification and is also useful for the identification of specific nucleic acid-protein interactions. A variety of other functional groups can be added to nucleic acids, such as sugars or glycans, amino acids, magnetic beads, gold nanoparticles, etc. For example, it has been recently reported that some RNAs are glycosylated within cells. Our sulfinate modification protocol could be used to generate large quantities of glycosylated RNAs to probe this new class of modified RNAs and their biological function. Our method enables a more diverse set of modifications and can be easily scaled to the larger quantities required for the industrial production of RNA therapeutics. It should be emphasized that the significance of this work lies in the ability to add carbon-carbon bonds to the nucleobases of RNA under mild, aqueous conditions and yet still retain an intact RNA backbone.

## Acknowledgments

We would like to thank Professor Stan Opella, Sang Ho Park and Anthony Mrse for their help with fluorine-19 NMR evaluation and Professor Yitzak Tor and Yao Li for their help with LC-MS experiments. We would also like to thank Tim Wiryaman for his preliminary experiments with PSMS-Zn modification of RNA. We also thank Phil Baran for helpful comments on the manuscript. This work was supported by the National Institute of General Medical Sciences of the National Institutes of Health (NIH) under award numbers R01GM123275 and R35GM141706 awarded to N.T. This work was also supported by the Howard Hughes Medical Institute (HHMI) Gilliam Fellowship awarded to A.H.

## Author Contributions

A.H., T.B., J.C.V., L.G. and N.T. designed the experiments. A.H., T.B., J.C.V., and L.G. performed the experiments. A.H., T.B., and N.T. wrote the manuscript with input from all authors.

## Methods

### In vitro transcription of GFP mRNA

Transcription reactions contained the following components: 40 μg linearized plasmid DNA, 40 mM Tris HCl pH 7.5, 25 mM MgCl2, 2 mM spermidine, 2.5 mM NTPs, 0.05% Triton X-100, 5 mM dithiothreitol, 5 μL T7 polymerase, 0.05 units of inorganic pyrophosphatase and water to a 1 mL final volume. Transcription reactions were run at 37 °C for a minimum of 3 hours and reactions were stopped with addition of 12 μL of 0.1 M CaCl2 and 20 units of TURBO DNase. Incubation with DNase proceeded for approximately 40 minutes at 37 °C. 6 units of Proteinase K was added to reactions and the reactions were incubated at 37 °C for an hour. Transcription reaction tubes were spun down for 5 minutes at 21,130 RCF and the supernatant was filtered through a 0.2 μm membrane. The RNA was purified by buffer exchange in centrifugal filters.

### Modification of nucleic acids with DAAS-Na

Reagents were added in the following order: (1) water (an amount should be added such that all other reagents reach their appropriate final concentrations after TBHP has been added) (2) appropriate buffer stock solution (we recommend sodium cacodylate, pH 6.5) (3) DAAS-Na stock solution, prepared in water (4) RNA stock solution, prepared in water (5) TBHP stock solution, prepared in water (ensure that the reaction is well-mixed immediately after TBHP addition). DAAS-Na final concentrations can be varied between 7.5-17.5 mM to alter labeling levels as shown in the “Results” section. However, whatever final concentration is chosen for DAAS-Na, ensure that TBHP remains at a 1.5-fold molar excess above DAAS-Na. We did not find that the reaction needed to be cooled prior to TBHP addition (as is often recommended in other sulfinate labeling protocols for small molecules), as our TBHP stock solution was already somewhat dilute (300 mM in water). Furthermore, the small reaction scales (~100 μL) ensured that there was no significant heat production upon addition of TBHP. When not being used on the benchtop, we always kept the TBHP stock solution at 4 °C. Final buffer concentrations are somewhat forgiving, though we recommend following our protocol of using sodium cacodylate, pH 6.5, at a 2.5-fold molar excess above DAAS-Na. We recommend avoiding MES and other similar buffers like MOPS and HEPES due to negative effects on yields^24,25^. However, even with a more well-behaved buffer like sodium cacodylate, using far more than necessary will also affect yields. RNA should be kept at a final concentration of 1 mg/mL, regardless of the concentration of DAAS-Na used. Once the reaction has been prepared and mixed thoroughly to ensure a homogenous distribution of reagents, it can be allowed to react at room temperature for ~24 hours.

Note on DAAS-Na concentrations above 17.5 mM: We observed that DAAS-Na concentrations above 17.5 mM tended to make the RNA precipitate during the course of labeling, with larger amounts of precipitate being formed as the DAAS-Na concentration was further increased. Considering that the DAAS label is quite hydrophobic, it is likely that labeling the RNA above a critical threshold causes it to no longer be significantly soluble in water, hence the precipitation. We observed that adding DMSO (to a final concentration of ~50%) helped dissolve the precipitate, while adding water had no effect, acting as further evidence that the RNA precipitated as a direct result of hydrophobicity due to high DAAS labeling.

### Purification of DAAS-RNA

Once the DAAS-Na labeling reactions were complete, purification of the resulting DAAS-RNA was relatively simple. The reaction mixture in its entirety was applied to a molecular weight cutoff filter of the appropriate cutoff size, such that all small molecules associated with the labeling reaction were capable of passing through the filter, leaving the DAAS-RNA behind. Enough filtration cycles were performed until sufficient purity of the DAAS-RNA was achieved. Water was used for buffer exchange, though any desired buffer can also be used.

### DBCO labeling of DAAS-RNA (Strain-promoted azide-alkyne cycloaddition – SPAAC)

Reagents were added in the following order: (1) water (an amount should be added such that all other reagents reach their appropriate final concentrations after the DBCO-labeled compound has been added) (2) appropriate buffer stock solution (we recommend sodium cacodylate, pH 6.5) (3) DAAS-RNA, in water (4) DBCO-labeled compound, prepared in water (ensure that the reaction is well-mixed after addition of the DBCO-labeled compound). The type and final concentration of buffer used is highly forgiving since the defining characteristic of biorthogonal chemistry is its ability to tolerate most common functionalities. We typically used 50 mM sodium cacodylate, pH 6.5, for DBCO reactions. DAAS-RNA and DBCO-labeled compound concentrations are also highly flexible; we used 0.5 mg/mL DAAS-RNA and ~500 μM DBCO-labeled compound. Ensure that enough DBCO-labeled compound is added for the expected amount of DAAS labels present on the DAAS-RNA, so that conjugation of every azide tag can be achieved. Once the reaction has been prepared and mixed thoroughly to ensure a homogenous distribution of reagents, it can be allowed to react at room temperature (or elevated temperature to ensure completion is reached) for ~24 hours. A small amount of EDTA can be added along with buffer, especially if the reaction is to be run at elevated temperature, in order to reduce any minimal backbone cleavage from trace metals.

### Purification of DBCO-DAAS-RNA

This procedure is identical to the “Purification of DAAS-RNA” procedure.

### Modification of nucleic acids with TFMS-Zn, TFMS-Na, and other sulfinate salts

This procedure is essentially identical to the “Modification of nucleic acids with DAAS-Na” procedure, aside from replacement of DAAS-Na with the sulfinate salt of choice. Note that zinc sulfinate salts tend to result in additional degradation beyond their sodium salt counterparts due to the ability of multivalent transition metals to catalyze phosphodiester backbone cleavage. Though zinc sulfinate salts have become popular due to their higher labeling yields, in general, we have found that the lower yields afforded by the sodium salts are more than compensated for by their lack of metal-catalyzed backbone cleavage, especially considering that the yield issue can be solved by simply increasing the concentration of the sodium sulfinate salt since it is relatively cheap compared to the RNA itself.

### ^19^F NMR sample preparation

The labeled RNA of interest was first digested with nuclease P1 according to provided protocols (water + 10x P1 reaction buffer + labeled RNA + nuclease P1). An excess of nuclease P1 above the manufacturer’s recommendation was added, along with an overnight incubation, to ensure complete digestion of the labeled RNA. After completion, directly to the nuclease P1 digest was added a known aliquot of ^19^F NMR internal standard (optional), along with D2O to approximately 33% D2O by volume. This entire mixture was then added to an NMR tube for analysis.

### LC/MS analysis of labeled DNA and RNA samples

Labeled DNA/RNA oligos were analyzed on a Waters I-Class LC with a Waters BEH C18 column (2.1×55 mm, 1.7 μm, 130 Å) using a gradient of 114 mM HFIP and 14 mM Et3N in water (A) and methanol (B) (0.3 mL/min, 10-26 %B over 10 minutes) at 60 °C. Identities were determined by ESI+.

## References

1. Frye, M., Harada, B.T., Behm, M. & He, C. RNA modifications modulate gene expression during development. Science 361, 1346–1349 (2018).

2. Kariko, K. et al. Incorporation of pseudouridine into mRNA yields superior nonimmunogenic vector with increased translational capacity and biological stability. Mol Ther 16, 1833–40 (2008).

3. Mulligan, M.J. et al. Phase I/II study of COVID-19 RNA vaccine BNT162b1 in adults. Nature 586, 589–593 (2020).

4. Yue, Y., Liu, J. & He, C. RNA N6-methyladenosine methylation in post-transcriptional gene expression regulation. Genes Dev 29, 1343–55 (2015).

5. Arango, D. et al. Acetylation of Cytidine in mRNA Promotes Translation Efficiency. Cell 175, 1872–1886.e24 (2018).

6. Ziehler, W.A. & Engelke, D.R. Probing RNA structure with chemical reagents and enzymes. Curr Protoc Nucleic Acid Chem Chapter 6, Unit 6.1 (2001).

7. Sawant, A.A. et al. A versatile toolbox for posttranscriptional chemical labeling and imaging of RNA. Nucleic Acids Res 44, e16 (2016).

8. Liu, Y., Sousa, R. & Wang, Y.X. Specific labeling: An effective tool to explore the RNA world. Bioessays 38, 192–200 (2016).

9. Flamme, M., McKenzie, L.K., Sarac, I. & Hollenstein, M. Chemical methods for the modification of RNA. Methods 161, 64–82 (2019).

10. Flood, D.T. et al. Synthetic Elaboration of Native DNA by RASS (SENDR). ACS Cent Sci 6, 1789–1799 (2020).

11. Gillingham, D. & Rasale, D. Direct and Selective Modification of RNA – An Open Challenge in Nucleic Acid Chemistry. Chimia (Aarau) 72, 777–781 (2018).

12. George, J.T. & Srivatsan, S.G. Posttranscriptional chemical labeling of RNA by using bioorthogonal chemistry. Methods 120, 28–38 (2017).

13. Trepotec, Z., Lichtenegger, E., Plank, C., Aneja, M.K. & Rudolph, C. Delivery of mRNA Therapeutics for the Treatment of Hepatic Diseases. Mol Ther 27, 794–802 (2019).

14. Ni, S. et al. Chemical Modifications of Nucleic Acid Aptamers for Therapeutic Purposes. Int J Mol Sci 18 (2017).

15. Komarova, N. & Kuznetsov, A. Inside the Black Box: What Makes SELEX Better? Molecules 24 (2019).

16. Gemmill, D., D’souza, S., Meier-Stephenson, V. & Patel, T.R. Current approaches for RNA-labelling to identify RNA-binding proteins. Biochem Cell Biol 98, 31–41 (2020).

17. Fujiwara, Y. et al. Practical and innate carbon-hydrogen functionalization of heterocycles. Nature 492, 95–9 (2012).

18. Zhou, Q. et al. Bioconjugation by native chemical tagging of C-H bonds. J Am Chem Soc 135, 12994–7 (2013).

19. Gui, J. et al. C-H methylation of heteroarenes inspired by radical SAM methyl transferase. J Am Chem Soc 136, 4853–6 (2014).

20. Ji, Y. et al. Innate C-H trifluoromethylation of heterocycles. Proc Natl Acad Sci U S A 108, 14411–5 (2011).

21. Langlois, B., Laurent, E. & Roidot, N. Trifluoromethylation of aromatic compounds with sodium trifluoromethanesulfinate under oxidative conditions. Tetrahedron Lett. 32, 7525–7528 (1991).

22. Ingle, S., Azad, R.N., Jain, S.S. & Tullius, T.D. Chemical probing of RNA with the hydroxyl radical at single-atom resolution. Nucleic Acids Res 42, 12758–67 (2014).

23. Agard, N.J., Prescher, J.A. & Bertozzi, C.R. A strain-promoted [3 + 2] azide-alkyne cycloaddition for covalent modification of biomolecules in living systems. J Am Chem Soc 126, 15046–7 (2004).

24. Baker, C.J. et al. Interference by Mes [2-(4-morpholino)ethanesulfonic acid] and related buffers with phenolic oxidation by peroxidase. Free Radic Biol Med 43, 1322–7 (2007).

25. Ichiishi, N. et al. Protecting group free radical C-H trifluoromethylation of peptides. Chem Sci 9, 4168–4175 (2018).

