## Supplementary Information for "Diverse chemical functionalization of nucleobases within long RNAs using sulfinate salts"

| Labeling Reaction | GFP mRNA | Sodium Phosphate pH 6.8 | TFMS-Na | TBHP |
|---|---|---|---|---|
| B1 | 1 mg/mL | 25 mM | 0 mM | 15 mM |
| B2 | 1 mg/mL | 25 mM | 20 mM | 15 mM |

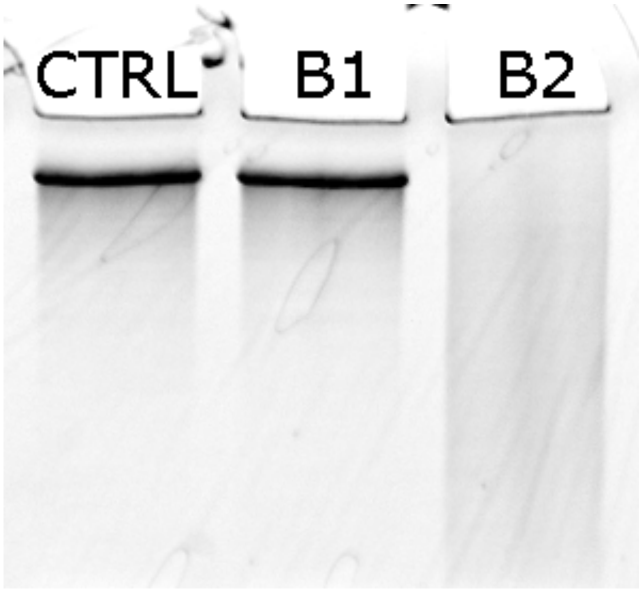


Supplementary Figure 1 (above): Final concentrations of all reagents are shown for reactions B1 and B2. All reactions were in aqueous solution, at room temperature, and run for 22 hours. Denaturing PAGE of the recovered RNA from reactions B1 and B2 (lanes 2 and 3) is shown along with unlabeled RNA as a control (lane 1). This gel was stained with ethidium bromide to make the RNA visible.

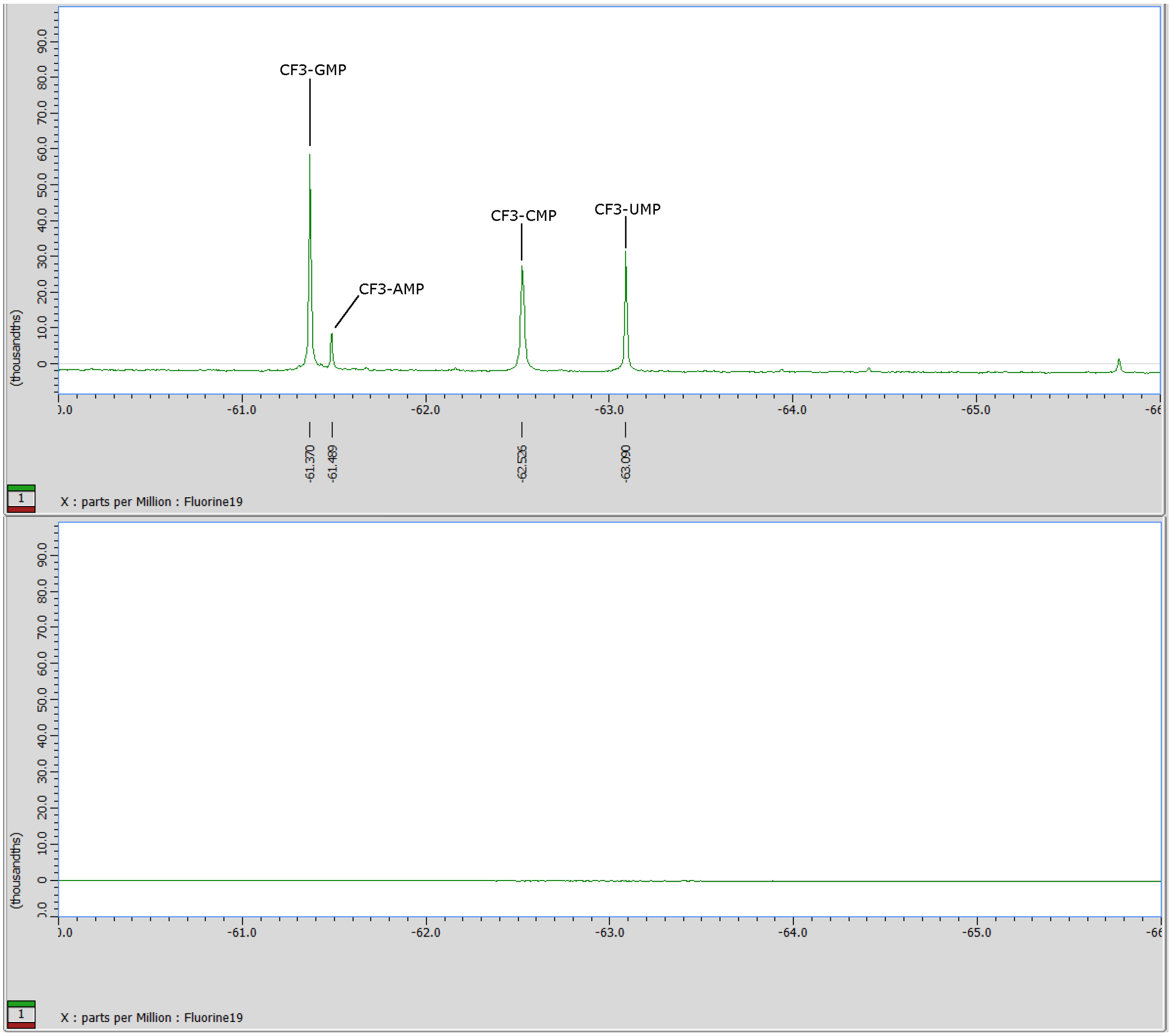


Supplementary Figure 2 (above): The top $^{19}F$ NMR spectrum corresponds to the nuclease P1 digest of the $CF_3$-labeled RNA from reaction B2. Peak identities were determined by labeling a sample of each pure nucleoside 5'-monophosphate individually, followed by analysis of its chemical shift. These chemical shifts correspond exactly to those observed with the nuclease P1 digest of the $CF_3$ labeled RNA since this particular nuclease yields nucleoside 5'-monophosphates. The bottom $^{19}F$ NMR spectrum corresponds to the nuclease P1 digest of the RNA from reaction C4, which turned out to be unlabeled, as the lack of peaks suggests.

| Labeling Reaction | GFP mRNA | Sodium Cacodylate, pH 6.5 | Sodium Acetate, pH 5 | Tris-HCl, pH 7.5 | TFMS-Zn | TBHP | TBHP Addition Condition |
|---|---|---|---|---|---|---|---|
| C1 | 0.161 mg/mL | 25 mM | 0 mM | 0 mM | 1 mM | 2.5 mM | No cooling before or after TBHP addition |
| C2 | 0.161 mg/mL | 25 mM | 0 mM | 0 mM | 1 mM | 2.5 mM | Cooling before and after TBHP addition |
| C3 | 0.161 mg/mL | 0 mM | 25 mM | 0 mM | 1 mM | 2.5 mM | Cooling before and after TBHP addition |
| C4 | 0.161 mg/mL | 0 mM | 100 mM | 0 mM | 1 mM | 2.5 mM | Cooling before and after TBHP addition |
| C5 | 0.161 mg/mL | 0 mM | 0 mM | 25 mM | 1 mM | 2.5 mM | Cooling before and after TBHP addition |
| C6 | 0.161 mg/mL | 25 mM | 0 mM | 0 mM | 1 mM | 2.5 mM | TBHP addition spread over 2 hours, cooling before and after each TBHP addition |

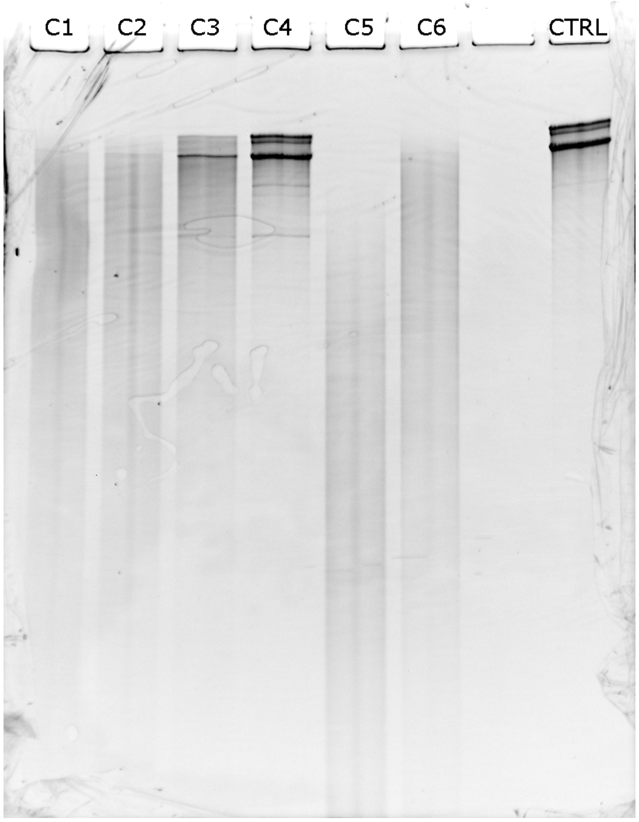


Supplementary Figure 3 (above): Final concentrations of all reagents are shown for reactions C1-C6. All reactions were in aqueous solution, at room temperature. Reactions with “cooling before and after TBHP addition” were cooled to 0 °C in ice-water baths (in order to test the effect of cooling on the reaction – later analysis showed that this made no difference). All reactions were run for 19 hours. Denaturing PAGE of the recovered RNA from reactions C1-C6 (lanes 1-6) is shown along with unlabeled RNA as a control (lane 8). This gel was stained with ethidium bromide to make the RNA visible. Note that the in vitro transcription reaction which produced this RNA also produced longer side products due to incomplete restriction digestion of the template plasmid. However, this doesn’t affect the overall conclusions that can be drawn from the gel regarding RNA degradation.

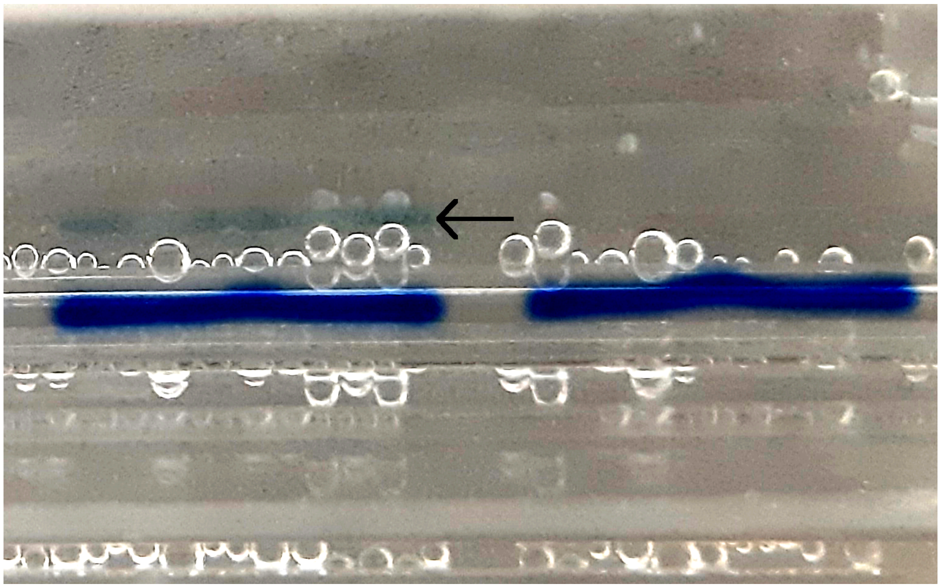

Supplementary Figure 4 (above, contrast enhanced): The band for DBCO-DAAS-RNA A5 is shown on the left (with the arrow pointing towards it) just a few minutes after the denaturing PAGE run was started. Sample A6 to the right is not visible because it is not labeled with dye. Note that the dark blue bands are the bromophenol blue loading dye, not the RNA.

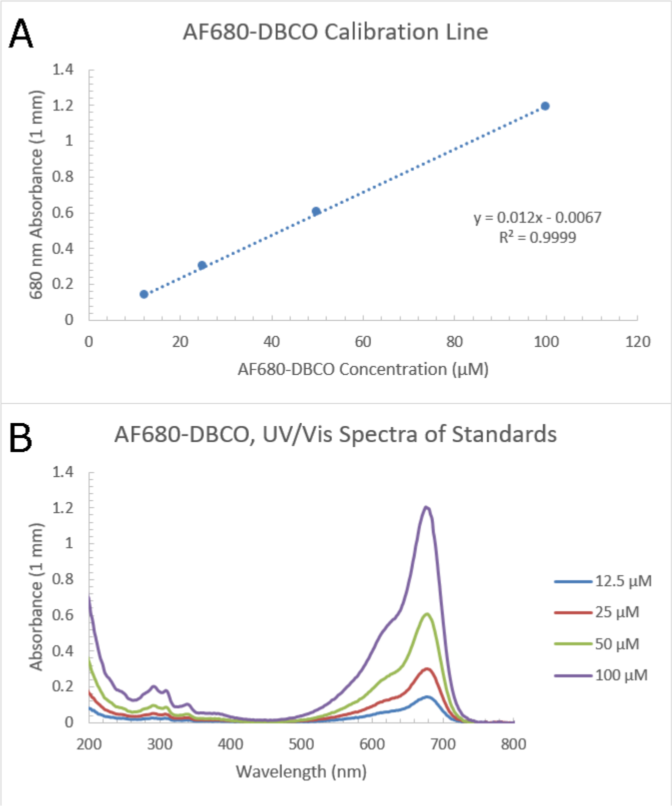


Supplementary Figure 5 (above): The calibration line (Panel A) and UV/Vis absorbance spectra (Panel B) for the four AF680-DBCO standard solutions are shown.

| $^{19}F$ NMR Analysis of $CF_3$ Labeling Percentages for Reaction B2 | |
|---|---|
| GMP | 20% |
| AMP | 3% |
| CMP | 14% |
| UMP | 18% |

Supplementary Table 1 (above): $CF_3$ labeling percentages for reaction B2, derived by $^{19}F$ NMR peak area analysis using a sodium trifluoroacetate internal standard. Percentages are defined relative to the total amount of each nucleotide that was originally present in the RNA before nuclease P1 digestion. All labeling percentages shown here are derived from the corresponding peaks in Supplementary Figure 2.

| DAAS-Na Concentration (mM) | % Labeling |
|---|---|
| 0 | 0.0 |
| 7.5 | 0.3 |
| 10.0 | 0.9 |
| 12.5 | 1.7 |
| 15.0 | 3.1 |
| 17.5 | 4.2 |

Supplementary Table 2 (above): The % labeling vs. DAAS-Na concentration used in each corresponding labeling reaction is shown.
